# Encoding-related fMRI BOLD activity predicts subsequent memory for studied scenes, but not subsequent identification of perceptually similar lures

**DOI:** 10.64898/2026.06.17.732733

**Authors:** Ayse N.Z. Aktas, Sabina Srokova, Michael D. Rugg

## Abstract

Pattern separation is widely regarded as a hippocampally mediated process that reduces interference between memories of similar experiences. Performance on the Mnemonic Similarity Task (MST), where the requirement is to discriminate between studied items (targets) and perceptually similar lures, is commonly held to depend on pattern separation. Specifically, it has been proposed that similar lure identification is supported by a recall-to-reject strategy, whereby lures are identified as a result of the retrieval of the corresponding studied item. According to this proposal, therefore, the encoding operations that support successful target recollection and successful lure identification should be closely similar, since both mnemonic judgments depend upon subsequent recollection of the study item. Here, using fMRI, we examined this prediction. In samples of cognitively healthy young and older adults, we employed a three-choice MST (target/lure/new) with scene and object images as test items. Using ROI-based univariate and multivoxel analyses, we assessed whether encoding-related activity was predictive of the identification of target and similar lure items on the subsequent memory test. The activity elicited by scene images predicted memory performance for subsequently presented targets but not for corresponding similar lures, contrary to the recall-to-reject hypothesis. No effects could be identified for either class of object test items. The magnitude of the encoding effects for the scene targets was age-invariant, and, moreover, the univariate scene SMEs demonstrated a robust, age-invariant association with the target-lure discriminability metric.

**Highlights:**

- Encoding fMRI activity predicted scene recognition but not lure discrimination.
- Encoding-related fMRI effects showed no age-related differences.
- Encoding fMRI effects for scenes predicted mnemonic discrimination performance.
- No encoding-related fMRI effects were detected for objects.

## 1. Introduction

Episodic memory, the remembrance of unique events (Tulving, 1972), relies on the ability to minimize interference from other encoded events that have overlapping features. Healthy aging is associated with declines in episodic memory (Craik & Salthouse, 2000), and memory difficulties represent the most common cognitive complaint in older adults (Mol et al., 2007).

A core process thought to support episodic memory, especially under conditions of interference, is pattern separation. Pattern separation supports the formation of decorrelated representations of similar inputs, thereby reducing their potential for representational overlap (O’Reilly & McClelland, 1994). Beginning with Marr (1971), the process of pattern separation has been linked to the mammalian hippocampus (Leutgeb et al., 2007; Bakker et al., 2008; Neunuebel & Knierim, 2014; Sakon & Suzuki, 2019; but see Amer & Davachi, 2023).

The widely used Mnemonic Similarity Task (MST), a modified recognition memory test, is believed to tax hippocampally-mediated pattern separation processes (Stark et al., 2023). In the MST, participants study a series of items and, at test, encounter exact repetitions of studied items (hereafter referred to as *targets*) intermixed with perceptually *similar lures* (and, in some versions of the task, perceptually dissimilar items also). It has been argued that successful identification of similar lure items depends heavily on a ‘recall-to-reject’ strategy, such that rejecting similar lures as targets requires the retrieval (recollection) of the corresponding studied item, which is then compared with the lure (Norman & O’Reilly, 2003; Yassa & Stark, 2011; Kim & Yassa, 2013; Lee and Stark, 2023; Parks et al., 2026). Thus, for recall-to-reject to succeed, the originally studied item must be encoded and subsequently retrieved with sufficient fidelity to allow correct rejection of its similar lure (Kirwan & Stark, 2007).

The MST is frequently employed in aging studies, when older adults typically show worse performance than younger participants (Stark et al., 2015; Stark & Stark, 2017; Reagh et al., 2018). This age difference in mnemonic discrimination has been interpreted as evidence for an age-related decline in hippocampal pattern separation (Yassa et al., 2011; Stark et al., 2019). Furthermore, relative to cognitively healthy controls, low performance on the MST has been reported for individuals diagnosed with Alzheimer’s Disease (AD) (Ally et al., 2013) and with mild cognitive impairment (MCI) (Stark et al., 2013), and MST performance has been proposed as a marker of early AD (Ally et al., 2013; Vanderlip et al., 2024). The importance of examining the mechanisms underlying performance on the MST arises from such interpretations and its now widespread use.

Examining the processes involved in forming memory representations at encoding can arguably provide insight into the neurocognitive mechanisms supporting later memory performance on the MST. According to the ‘recall-to-reject’ accounts (see above), correct target endorsements and correct lure rejections should both rely largely on recollection-based processes, such that failure to encode an item with sufficient fidelity should increase the likelihood of falsely endorsing a lure as a target at test. Consistent with this proposal, eye-tracking evidence indicates that study items whose corresponding lures were later falsely endorsed as targets attracted fewer fixations during encoding, whereas fixation counts did not differ between items later associated with correct target recognition and correct lure rejection (Molitor et al., 2014).

Replicating this general pattern, Bjornn et al. (2022) reported that the number of fixations directed toward a study item was inversely related to the likelihood of a false alarm to its corresponding lure, underscoring the importance of encoding highly distinctive representations. Extending these findings, Walheim et al. (2023) reported that accurate target recognition and lure rejection were both associated with greater on-task attention during encoding, suggesting that attentional engagement supports the formation of higher-fidelity memory representations.

In light of these considerations, we examined encoding-related neural activity by examining functional magnetic imaging (fMRI) subsequent memory effects (SMEs), which index the extent to which fMRI BOLD responses elicited during encoding differ as a function of later memory outcome. In the case of univariate SMEs, positive SME effects (relatively greater BOLD activity for items that go on to receive an accurate rather than an inaccurate memory judgment) have been consistently associated with subsequent recollection of a study item across a wide range of stimulus materials and neural regions (Cansino et al., 2002; Davachi et al., 2003; Ranganath et al., 2004; Uncapher & Rugg, 2005, Preston et al., 2009; see Kim, 2011 for review). In contrast, SMEs linked to familiarity-based memory judgments tend to be more regionally restricted and less robust (Davachi et al., 2003; Ranganath et al., 2004; Uncapher & Rugg, 2005; 2008). Prior findings suggest that positive SMEs do not differ dramatically as a function of age, although quantitative differences may be present (e.g., Brehmer et al., 2016; de Chastelaine et al., 2016; Liu et al., 2021; see Maillet & Rajah, 2014 for review). However, it remains unclear whether younger and older adults rely on similar encoding strategies for targets and perceptually similar lures in the context of the MST. Moreover, SMEs predicting accurate vs. inaccurate lure judgments have not previously been examined, even in young adults, leaving open the question of whether SMEs differ as a function of subsequent target versus lure detection.

In the present study, we quantified univariate subsequent memory effects (SMEs) in a priori regions of interest (ROIs) using a scaled metric, computed as the difference in mean encoding-related BOLD activity between memory outcomes normalized by the pooled trial-wise standard deviation (akin to Cohen’s *d*). This metric provided an effect size measure that facilitates comparison across regions and groups. Additionally, we conducted an exploratory mass-univariate analysis to identify SMEs across the whole brain. Complementing these univariate analyses, we also employed ROI-based multivoxel pattern similarity analysis (PSA) to assess the similarity of voxel-wise activation patterns during encoding as a function of subsequent memory outcomes. This approach indexed the extent to which neural representations were more similar for within than between outcome conditions, providing a multivoxel analog of the SME (see Yu et al., 2025, for an example of a study employing a closely analogous approach). Together, these methods allowed us to test whether the magnitude or the patterning of encoding-related activity differed as a function of subsequent target and lure performance, and whether these effects varied between younger and older adults.

We examined univariate and multivoxel SMEs associated with correct target endorsements and correct lure detection in a three-choice version of the MST. The experimental stimuli consisted of object and scene images, allowing examination of these metrics in category-selective occipitotemporal cortex (see Grill-Spector & Weiner, 2014, for review), specifically, the scene-selective parahippocampal place area (PPA; Epstein & Kanwisher, 1998) and the object-selective lateral occipital complex (LOC; Grill-Spector et al., 2001). Participants, who comprised samples of younger and older adults, underwent fMRI during the encoding phase of the task. In a subsequent out-of-scanner memory test, they classified test items as ‘Old’ (exact repetitions from the study phase), ‘Similar’ (perceptually related lures corresponding to studied items), or ‘New’ (novel items unrelated to studied images). This design enabled us to directly relate encoding-related neural activity to subsequent memory outcomes. Critically, within this framework, if a recall-to-reject strategy supports accurate mnemonic discrimination, SMEs should be evident not only for study items correctly endorsed as targets, but also for items whose corresponding lure was correctly rejected as a target.

The study utilized data from an experiment first reported by Srokova et al. (2024) to address a different set of questions. Unlike in Srokova et al., here we focus on those items presented during the in-scanner encoding phase that were subsequently tested in an out-of-scanner memory task, items that were excluded from the analyses described in Srokova et al. (2024). This approach allowed us to directly examine encoding-related neural activity as a function of subsequent memory performance, providing a novel examination of subsequent memory effects at encoding.

## 2. Methods

The dataset employed in this manuscript was first described by Srokova et al. (2024). The experimental materials, experimental and MRI methods, and the behavioral results are described in detail in that paper and summarized here for the convenience of the reader. The analysis of the fMRI SMEs that are the focus of the present study has not been reported previously.

### 2.1. Participants

Twenty-three young adults (18-30 years; M = 21.6; SD = 3.5) and 24 older adults (65-75 years; M = 68.8; SD = 3.8) were included in the present analyses. All participants were right-handed, had normal or corrected-to-normal vision, and were fluent English speakers. Exclusion criteria included a history of neurological or psychiatric disease, substance abuse, diabetes, or current or recent use of prescription medication affecting the central nervous system.

Before the MRI session, all participants underwent a comprehensive neuropsychological test battery. Inclusion and exclusion criteria were selected to minimize the likelihood of including older participants with mild cognitive impairment or early dementia (see Srokova et al., 2024, for neuropsychological test results).

### 2.2. fMRI Data Acquisition and Preprocessing

MRI data were acquired at the Sammons BrainHealth Imaging Center at the University of Texas at Dallas using a Siemens Prisma 3 T scanner equipped with a 32-channel head coil. Functional images were acquired with a T2*-weighted BOLD-sensitive echoplanar imaging (EPI) sequence (flip angle = 70°; FOV = 220 × 220 mm; voxel size = 2 × 2 × 2 mm; TR = 1.52 ms; TE = 30 ms; 66 slices, multiband factor = 3). A 3D MPRAGE pulse sequence was employed to acquire a T1-weighted whole-brain anatomical scan (FOV = 256 × 256 mm; voxel size = 1 × 1 × 1 mm; 160 slices; sagittal acquisition). Following the last imaging run, a dual-echo fieldmap sequence, matching the EPI sequence’s 3D characteristics, was acquired with TEs of 4.92 and 7.38 ms, producing two magnitude images and a pre-subtracted phase image.

MRI data were preprocessed using Statistical Parametric Mapping (SPM12, Wellcome Department of Cognitive Neurology) and custom MATLAB code (MathWorks). The functional data were preprocessed in six steps. First, we calculated voxel displacement maps prior to the fieldmap correction. Second, SPM’s realign and unwarp procedure was applied. Third, the functional images were reoriented along the anterior and posterior commissures, then spatially normalized to SPM’s EPI template, and renormalized to an age-unbiased sample-specific EPI template (de Chastelaine et al., 2011, 2016b). Lastly, the functional data were smoothed with a 5 mm FWHM Gaussian kernel.

### 2.3. Materials & Procedure

The experimental stimuli were displayed with PsychoPy v2021.1.3 (Peirce et al., 2019). The study phase took place within the MRI scanner, whereas the subsequent memory test occurred after participants exited the scanner.

Figure 1 illustrates the study and test tasks along with examples of experimental stimuli. At study, participants viewed 120 images of scenes and 120 images of objects. 144 images (72 per image category) went on to be tested in the post-scan retrieval task. The studied images that did not go on to be tested were part of the neural selectivity analysis reported by Srokova et al., (2024) and are not discussed in the present study. The study phase consisted of 9 scanner runs, each lasting 4 min 38 s. On each trial, participants were presented with a red fixation cross (500ms), followed by the study image (2s) and then a white fixation cross (2s). Participants had a total of 4 seconds after the image onset to make a subjective “indoor/outdoor” judgment regarding the presented scene or object. A total of 126 null trials, each consisting of a white fixation cross displayed at the center of the screen, were randomly distributed between the critical trials.

**Figure 1.**
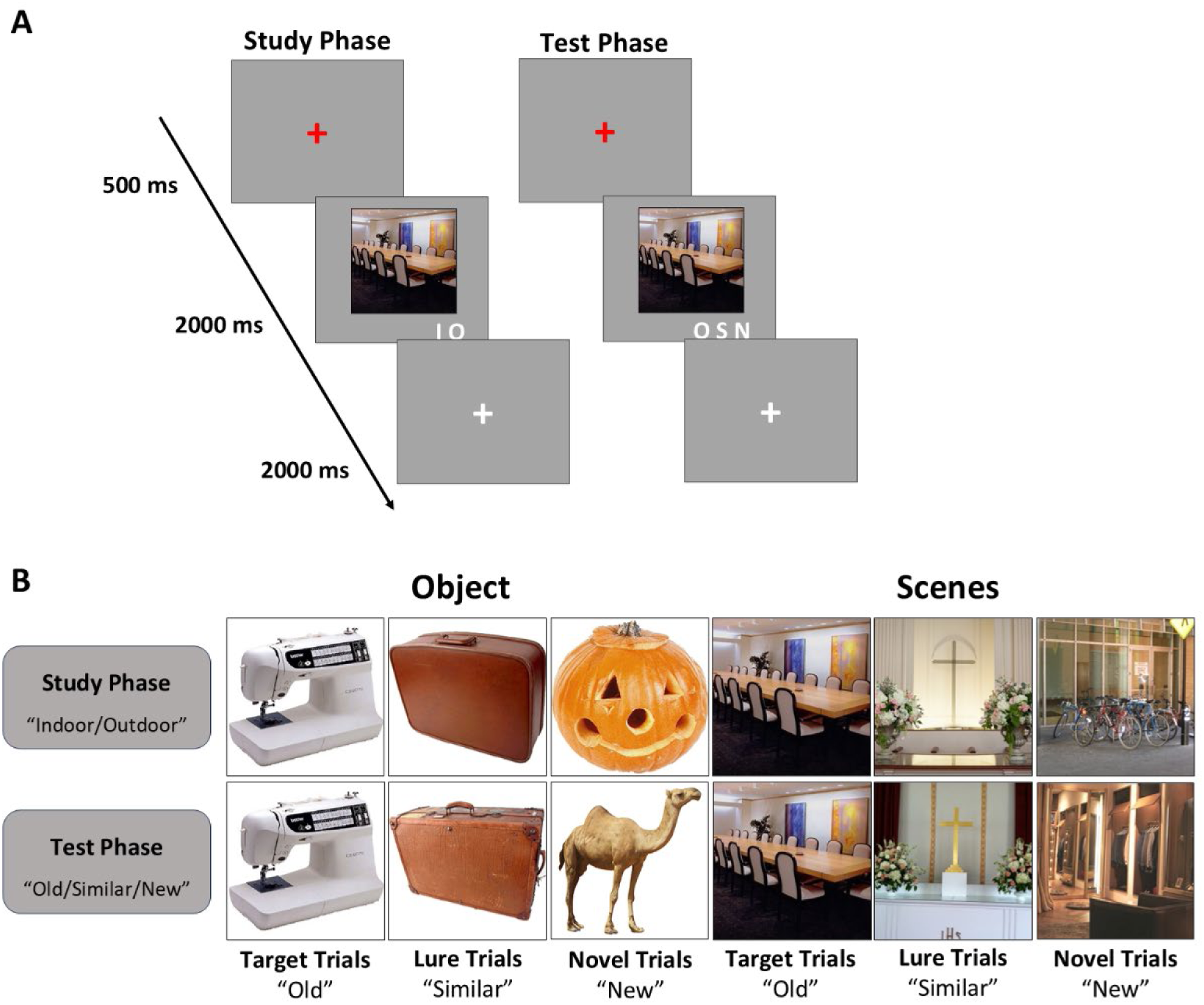
**(A)** Schematic of the study and test tasks. During the study phase, participants made ‘indoor/outdoor’ judgements, and at test, they made ‘old/similar/new’ judgements **(B)** Examples of target, lure, and new trials and corresponding correct responses for each of the two image categories (scenes and objects).

The test phase was completed outside of the MRI scanner. Participants viewed a total of 108 scene and 108 object images. For each image category, 36 trials were repetitions of images that the participant had viewed during the study phase (“targets”), 36 trials were images that were perceptually similar to previously viewed study images (“lures”), and 36 trials were presentations of new images. Each trial in the test phase started with a red fixation cross (500 ms), followed by the test image (2s), and concluded with a white fixation cross (2s), allowing for a 4-second response window. Participants were instructed to indicate whether the test image exactly matched one of those they had viewed during the study phase (“old”), was perceptually similar to an image from the study phase (“similar”), or was a completely new image (“new”). The stimuli were used to create 24 stimulus lists yoked between pairs of younger and older adults.

### 2.4. Behavioral Data Analysis

Behavioral responses from the subsequent memory test were separated according to item category (scenes and objects), trial type (target, lure, new), and participant endorsement (‘Old’, ‘Similar’, ‘New’) and binned separately by age group (older adult, younger adult). The respective proportions were then analyzed using a 2 (age) x 2 (category) x 3 (trial type) x 3 (endorsement) mixed effects ANOVA.

Additionally, item memory and ‘target-lure discriminability’ (TLD) indices were computed. Item memory was estimated as the difference between the proportion of old items correctly endorsed “Old” (item hits) and the proportion of new items incorrectly endorsed as “Old” (false alarms). For the reasons advanced by Srokova et al. (2024; see also Loiotile & Courtney, 2015), mnemonic discrimination was quantified with the TLD index, calculated as the difference between the proportion of old trials correctly endorsed “old” and the proportion of lure trials incorrectly endorsed “old.” The item and TLD metrics were entered into separate 2 (age) x 2 (category) mixed ANOVAs. Note that these behavioral metrics and their analyses were first reported by Srokova et al. (2024).

### 2.5. fMRI Analysis

The primary fMRI analyses were region of interest (ROI) based, focusing on the scene-selective parahippocampal place area (PPA) and the object-selective lateral occipital cortex (LOC). However, because SMEs are not restricted to canonical category-selective regions, we also report an exploratory whole-brain analysis.

#### 2.5.1. ROI Analysis ROI selection

As noted above, the two *a priori* ROIs were the PPA and LOC. The regions were defined by the intersection between category-selective univariate activity and anatomical labels provided by the Neuromorphometrics atlas (available in SPM12) (Koen et al., 2019; see also Srokova et al., 2020).

Before selecting the ROIs, the fMRI data were analyzed using a two-stage univariate GLM approach, following the procedures detailed by Srokova et al. (2024) and described below. To avoid circularity and enable an unbiased selection of the ROIs across the two age groups, a ‘leave-one-pair-out’ procedure was employed, following the approach outlined by Hill et al. (2021). This involved iteratively excluding a younger and older adult pair before estimating group-level GLMs and using the remaining sample data to identify the ROIs for the excluded pair. The parahippocampal and fusiform gyri were used to identify the parahippocampal place area (PPA) through a scene > object contrast (left PPA, M = 147 voxels; SD = 4.5; right PPA, M = 156 voxels; SD = 1.7), while the inferior and middle occipital gyri defined the lateral occipital complex (LOC) via an object > scene contrast (left LOC, M = 286 voxels; SD = 7.5; right LOC, M = 258 voxels; SD = 6.4). Both contrasts were height thresholded at p< .001 with a cluster extent threshold of p < 0.05 (FWE-corrected).

### Univariate SME metric applied within the ROIs

Separate first-level GLMs were constructed for each participant to obtain single-trial parameter estimates using a “least-squares-all” model (Rissman et al., 2004; Mumford et al., 2014) in which each trial was modeled by a separate 2-second boxcar regressor tracking image presentation. The six motion regressors reflecting rigid body translation and rotation were included as covariates of no interest. Trials were categorized based on subsequent memory responses into three types: (1) old items subsequently correctly endorsed as ‘old’ (target correct), (2) study items corresponding to a subsequently presented lure item that was correctly endorsed as ‘similar’ (lure correct), and (3) incorrect responses, which included any trials where the participant failed to accurately identify the target or lure. Because of the low number of item misses (incorrect ‘new’ endorsements), the incorrect trials were collapsed across all possible incorrect responses to targets (‘similar’ and ‘new’ endorsements) and lures (‘old’ and ‘new’ endorsements). These categories served as the basis for further contrasts to examine SMEs.

The single-trial data were employed to generate participant-wise subsequent memory metrics within each corresponding category-selective ROI, in the form of effect sizes (see Srokova et al., 2021 for a comparable approach in the case of metrics of neural selectivity and memory reinstatement). The metric for target trials (Target SME) was computed for each ROI as the difference between the mean fMRI BOLD response elicited by correctly recognized targets versus all encoding trials that went on to receive an incorrect response, divided by the pooled inter-trial standard deviation (SD; see equation 1 below). The SME metric for lures (Lure SME) was computed analogously for each ROI by computing the difference between the mean BOLD response for studied items corresponding to correctly identified lure items and the mean BOLD response for all study trials that went on to receive an incorrect response, divided by the pooled SD.

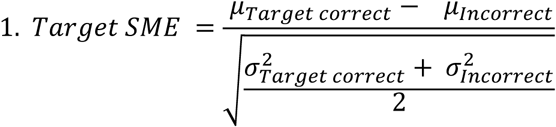

The SME metric was estimated for each participant within each of the four ROIs (left PPA, right PPA, left LOC, right LOC), separately for each image category. However, since the left and right ROIs showed no interaction with any other variable in preliminary analyses, they were combined into a single bilateral ROI (PPA and LOC), resulting in two metrics per participant: one for scenes in the PPA and one for objects in the LOC. Positive values signify greater BOLD activity for subsequently correctly endorsed trials than incorrectly endorsed (positive SMEs). Note that the scaling function of the SME metric renders it insensitive to individual variation in the gain of the hemodynamic transfer function (HRF), thus making it insensitive to the possibly confounding effects of group or individual differences in this variable (Liu et al., 2013).

### Multivoxel SME metrics: Pattern Similarity Analysis

Single-trial beta estimates, as defined above, were entered into a multivoxel pattern similarity analysis (PSA) to derive a multivoxel subsequent memory effect (MV-SME) index . MV-SME metrics, the multivoxel analog of univariate SMEs, were operationalized as the difference between within-condition and between-condition similarity of voxel-wise activation patterns, capturing pattern similarity as a function of subsequent memory outcomes (Figure 2). All similarity calculations were performed within image categories. To avoid confounding within-run autocorrelation effects (Mumford et al., 2014), all similarity metrics were calculated using correlations between trials from different scanner runs. For the target MV-SMEs, the metric was computed as the difference between the average within-condition similarity (i.e., the correlation between each study item that was later correctly recognized as a target with all other later correctly recognized targets) and the average between-condition similarity (i.e., the correlation between each study item that was correctly recognized as a target with all study items that later received an incorrect memory response). Similarly, lure MV-SMEs were defined as the difference between within-condition similarity (i.e., the correlation between each study item that was later correctly identified as a lure with all other correctly identified lures) and between-condition similarity (i.e., the correlation between each study item that was correctly identified as a lure with all study items that later received an incorrect memory response). Thus, higher MV-SME values signify greater neural similarity between study items that went on to be correctly identified as targets or whose lures were correctly identified.

**Figure 2.**
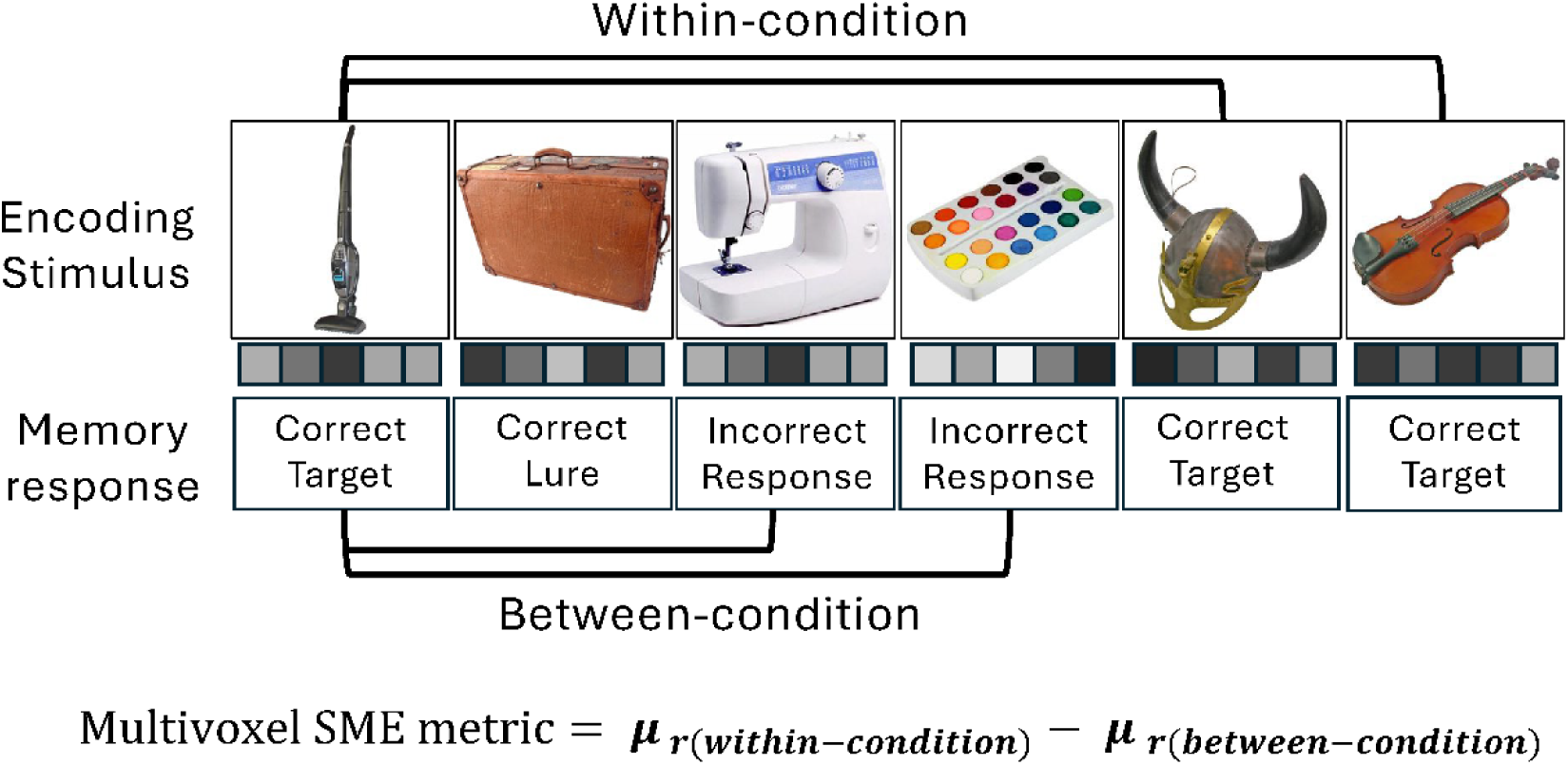
Schematic illustration of the multivoxel subsequent memory effect (MV-SME) calculation. MV-SME was computed as the difference between within-condition similarity (average correlation between each item and all other items with the same subsequent memory outcome) and between-condition similarity (average correlation between each item and all items with an incorrect subsequent memory outcome). Higher MV-SME values reflect greater neural similarity among items associated with successful subsequent memory performance.

#### 2.5.2. Whole-brain analysis

The functional MRI data were also analyzed with a two-level mass-univariate approach. In a series of first-level GLMs, whole-brain data for each participant were separately modeled to estimate neural activity elicited on each study trial. The trials were segregated by the memory judgment the study items attracted on the subsequent memory test, separately for scenes and objects, to create three events of interest for each item category: target correct (old items subsequently correctly endorsed as ‘old’), lure correct (study items corresponding to a subsequently presented lure item that was correctly endorsed as ‘similar’), and all items that received an incorrect judgement at test. As with the single-trial analyses described previously, the low number of item misses necessitated collapsing incorrect trials across erroneous responses to both targets and lures. Neural activity elicited by the 6 events of interest was modeled by convolving a canonical hemodynamic response function with a 2-second boxcar that onset at the time of presentation of the stimulus image. Additional regressors included a single column for trials of no interest (filler trials, no response trials, and items attracting responses occurring within 500ms post-stimulus onset), 6 motion regressors reflecting rigid-body translation and rotation, and spike covariates regressing out volumes with displacement greater than 1 mm or 1° in any direction. The fMRI time series from each scanning session was concatenated into a single time series using the spm_fmri_concatenate function.

To identify voxels that were sensitive to the subsequent memory judgment (target correct, lure correct, incorrect), the parameter estimates for each event of interest were taken forward to a second-level GLM. The GLM implemented a 2 (age) by 6 (event of interest) mixed factorial ANOVA (note that, as implemented in SPM, the ANOVA employed a single, pooled error term). All contrasts were tested against a height threshold of *p* < .001 (uncorrected) and a cluster extent threshold of *p* < .05, FWE corrected (*k* ≥ 75).

As with the single-trial data, SMEs were obtained by contrasting correctly identified targets or lures with incorrect trials, separately for scenes and objects. Thus, target SMEs were identified by a ‘Target Correct > Incorrect’ contrast, and lure SMEs were identified by a ‘Lure Correct > Incorrect’ contrast. Additionally, a direct comparison was made between correctly identified targets and correctly identified lures (target>lure contrast).

We also analyzed ‘negative’ target and negative lure SMEs in a similar manner. Negative SMEs reflect higher BOLD activity for later incorrectly endorsed than correctly endorsed study items (negative target SME: incorrect>target correct; negative lure SME: incorrect>lure correct).

These analyses did not identify any significant clusters at either the predetermined or a more liberal statistical threshold. Therefore, they are not discussed further.

### 2.6. Associations between encoding-related neural metrics and mnemonic discrimination performance

Multiple regression analyses were conducted to examine associations between encoding-related neural metrics, univariate and multivoxel SMEs, and mnemonic discrimination performance across participants. Mnemonic discrimination was quantified using the previously defined TLD index. In all models, age group and the interaction between age group and the neural metric of interest were included to assess potential moderation by age.

To control the family-wise error rate, p-values were adjusted using the Holm–Bonferroni procedure (Holm, 1979) across the four planned regression models within each neural metric family (SME and PSA models were corrected separately). Unless otherwise mentioned, all reported p-values survived correction.

### Univariate SME metrics and mnemonic discrimination performance

Single-trial SME metrics were estimated separately for target and lure trials within each category-selective ROI (PPA and LOC). Four separate regression models were constructed for each combination of SME metric (target SME, lure SME) and ROI. In each model, SME metrics were entered as predictors of TLD, along with age group and an interaction term (age group × SME) to evaluate whether age moderates the relationship between SME magnitude and mnemonic discrimination performance.

### Multivoxel SME metrics and mnemonic discrimination performance

A separate set of multiple regression analyses examined the relationship between the MV-SME metrics and mnemonic discrimination performance. MV-SME values were computed separately for correctly identified target and lure trials within each category-selective ROI. Four regression models were constructed, with target and lure MV-SME entered as predictors of TLD, along with age group and an interaction term (age group × MV-SME metric) to test for potential age-related moderation along with age group and an interaction term (age group × MV-SME metric) to test for age moderation.

### 2.7. Statistical Analysis

Statistical analyses were performed using R statistical software (v4.2.2.; R Core Team, 2022). Analyses of variance were performed using the afex package (Singmann et al., 2023), and degrees of freedom were corrected for nonsphericity with the Greenhouse–Geisser procedure (Greenhouse and Geisser, 1959) when necessary. *t* tests and multiple regression analyses were performed using the t.test and lm functions, respectively, in base R. Partial correlations were conducted using pcor.test in the ppcor package (Kim, 2015).

### 2.8 Outlier definition

Outlying data points were defined as 2.5 standard deviations from their respective sample means and are identified in each of the figures illustrated below. Analyses were conducted with and after the removal of participants contributing an outlying data point. Unless otherwise stated, the outcomes of the analyses were statistically equivalent.

## 3. Results

### 3.1. Behavioral Results

As noted above, the behavioral findings were previously reported by Srokova et al., (2024); they are re-reported here for the reader’s convenience. Figure 3 illustrates item memory and TLD metrics.

**Figure 3.**
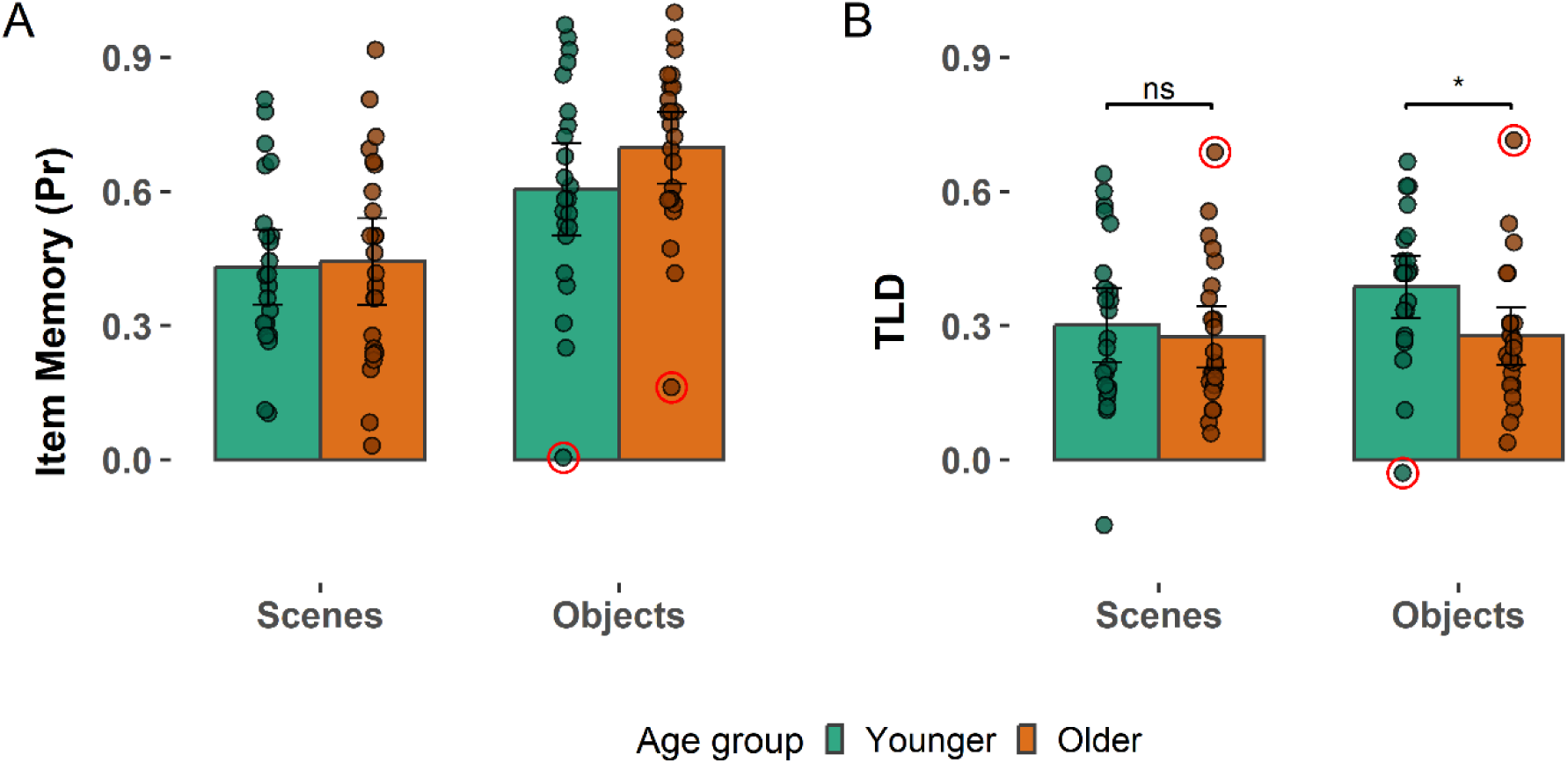
**(A)** Item recognition memory and **(B)** Target-lure discriminability (TLD) performance binned separately for younger and older adults. Error bars reflect 95% confidence intervals.

**Figure 4:**
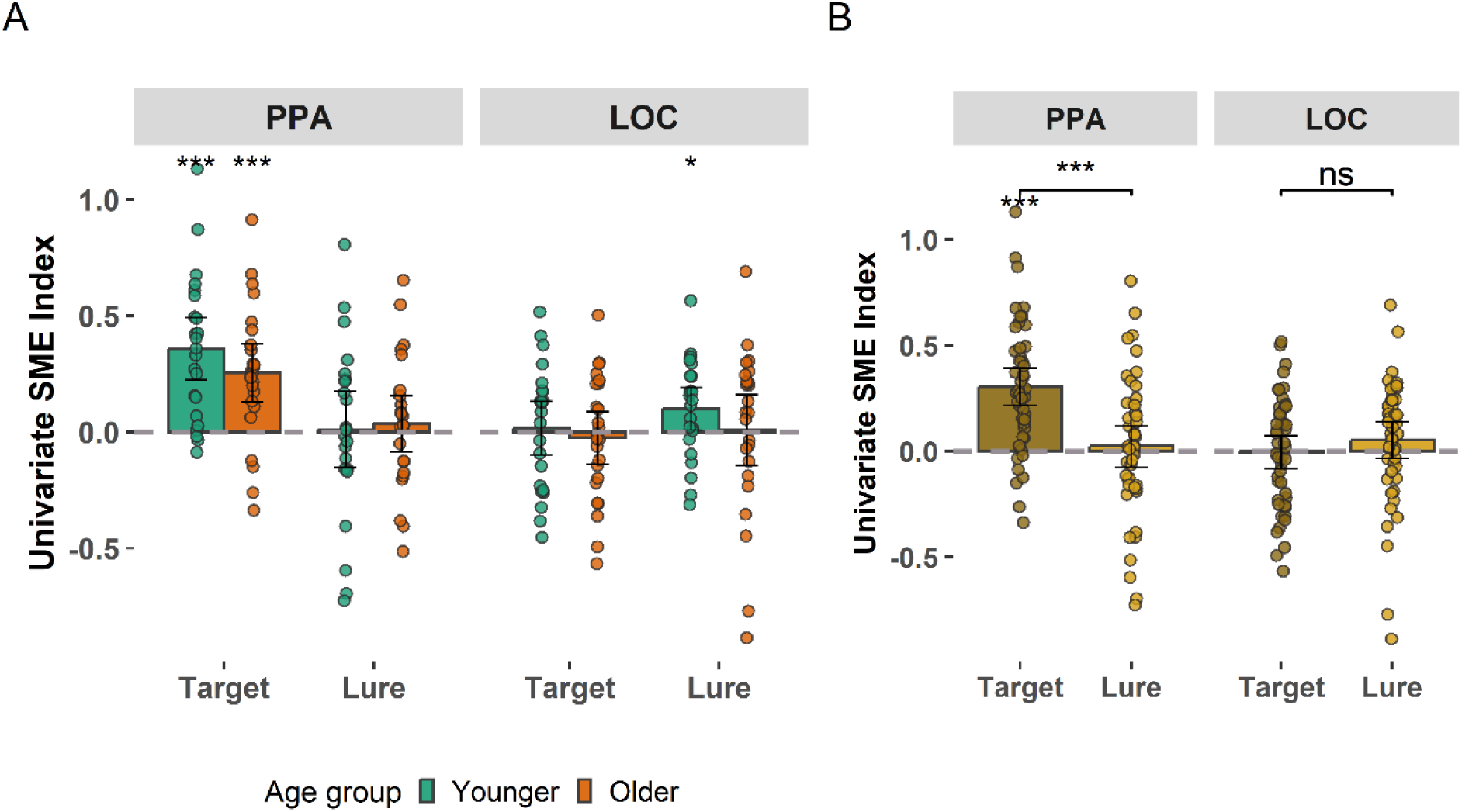
**(A)** Single-trial univariate SME metrics for younger and older adults, separately for scene images in the PPA and object images in the LOC. **(B)** SME metrics from the single-trial analysis collapsed across age group. Error bars represent 95% confidence intervals. Stars above individual bars indicate trial types where one-sample t-tests indicated the SMEs exceeded zero. The brackets indicate the outcomes of two-sample t-tests. (ns p > 0.05, *p < 0.05, **p < 0.01, ***p < 0.001).

#### Outlying data points are circled

The two behavioral metrics were subjected to separate 2 (age group) x 2 (image category) mixed effects ANOVAs. The ANOVA for item memory yielded a significant main effect of category (F(1, 45) = 73.027, p < 0.001, partial-η2 = 0.619), indicating better item memory for objects compared to scenes. However, neither the main effect of age group nor the age-by-category interaction reached significance (age: F(1, 45) = 0.852, p = 0.361, partial-η2 = 0.019; interaction: F(1, 45) = 2.482, p = 0.122, partial-η2 = 0.052). Consequently, there were no detectable age differences in item memory performance between young and older adults.

The ANOVA for the TLD metric revealed a significant main effect of category, *F*(1,45) = 4.74, *p* = .035, *partial η²* = .095, with higher performance for objects than scenes. The main effect of age group was not significant, F(1,45) = 2.34, p = .133, partial η² = .049; however, the age × category interaction reached significance, F(1,45) = 4.35, p = .043, partial η² = .088. Follow-up t-tests indicated that older adults exhibited lower discrimination performance than younger adults for objects, t(44.44) = 2.41, p = .020, but not for scenes, t(43.10) = 0.51, p = .613. Thus, the age-by-category interaction was driven by reduced object discrimination in older adults, with no age-related differences observed for scenes.

Exclusion of the outliers (circled in Figure 3B) resulted in a non-significant age group x category interaction (p = .053) while the main effect of age group became significant (p = .006); however, the main effect of category (p = .049) remained significant.

### 3.2. fMRI Results

The fMRI analyses examined fMRI BOLD activity associated with subsequent memory performance, focusing on SMEs and PSA derived from category-selective ROIs. Trial counts for each event type included in the analyses, across all conditions and age groups, are displayed in Table 1. Analyses were performed separately for the *a priori* scene- and object-selective ROIs and were complemented by whole-brain analyses to assess encoding-related effects beyond the predefined regions. Regression analyses were employed to assess whether these encoding-related metrics predicted, across participants, mnemonic discrimination performance.

**Table 1:** The mean (range) number of trials for events of interest: target correct, lure correct, and incorrect.

|  | Age group | Target correct | Lure correct | Incorrect |
| --- | --- | --- | --- | --- |
| Scenes | Younger | 16.7 (4-30) | 16.8 (10-26) | 37.5 (21-54) |
|  | Older | 18.9 (4-33) | 18.3 (10-27) | 34.2 (17-53) |
| Objects | Younger | 23.0 (3-35) | 18.7 (10-29) | 29.6 (15-55) |
|  | Older | 26.9 (15-36) | 12.1 (4-23) | 32.7 (12-47) |

#### 3.2.1. ROI Analyses Univariate SME metrics

Figure 4 illustrates the univariate SME metrics derived from the scene- and object-selective ROIs. The metrics were subjected to a 2 (age group) × 2 (ROI: PPA, LOC) × 2 (SME metric: target, lure) mixed-effects ANOVA. This resulted in a main effect of ROI (F(1,45) = 7.02, p = .011, partial η² = .135), a main effect of SME metric (F(1,45) = 11.36, p = .002, partial η² = .202), and a significant ROI × SME metric interaction (F(1,45) = 17.22, p < .001, partial η² = .277). The main effect of age group (F(1,45) = 1.30, p = .260, partial η² = .028), as well as the age group interactions with ROI (F(1,45) = 0.07, p = .793, partial η² = .002) and with the SME metric (F(1,45) = 0.35, p = .557, partial η² = .008) were not significant, and nor was the three-way interaction (F(1,45) = 1.19, p = .281, partial η² = .026).

To follow up the SME metric × ROI interaction, separate 2 (SME metric: target, lure) × 2 (age group) mixed-effects ANOVAs were conducted for the PPA and LOC (although age group did not interact with SME metric in the omnibus analysis, it was retained in the follow-up models given its central relevance to the study questions and to allow for the detection of potential age-related moderation within specific ROIs).

For the PPA, the ANOVA revealed a significant main effect for the SME metric (F(1,45) = 26.22, p < .001, partial η² = .368), no main effect of age group (F(1,45) = 0.27, p = .607, partial η² = .006), and no SME metric × age group interaction (F(1,45) = 1.36, p = .249, partial η² = .029). The main effect of SME metric reflected a larger target (M = 0.305, SD = 0.304) than lure SMEs (M = −0.005, SD = 0.266).

In contrast, the corresponding ANOVA for the LOC did not reveal any significant main effects or interactions (min. *p* = .194).

##### Multivoxel SME metrics

Figure 5 illustrates the MV-SME metrics separately for the scene-selective (PPA) and object-selective (LOC) ROIs. The data were analyzed using a 2 (Age Group) × 2 (ROI) × 2 (MV-SME metric: target, lure) mixed-effects ANOVA. This revealed a significant ROI × MV-SME metric interaction, F(1,45) = 19.23, p < .001, partial η² = .30. There were no significant main effects of MV-SME metric, F(1,45) = 3.74, p = .059, partial η² = .077, ROI F(1,45) = 1.29, p = .262, partial η² = .03, or age group, F(1,45) = 0.14, p = .715, partial η² < .01, and no other significant interactions involving age (min. *p* = .333).

**Figure 5.**
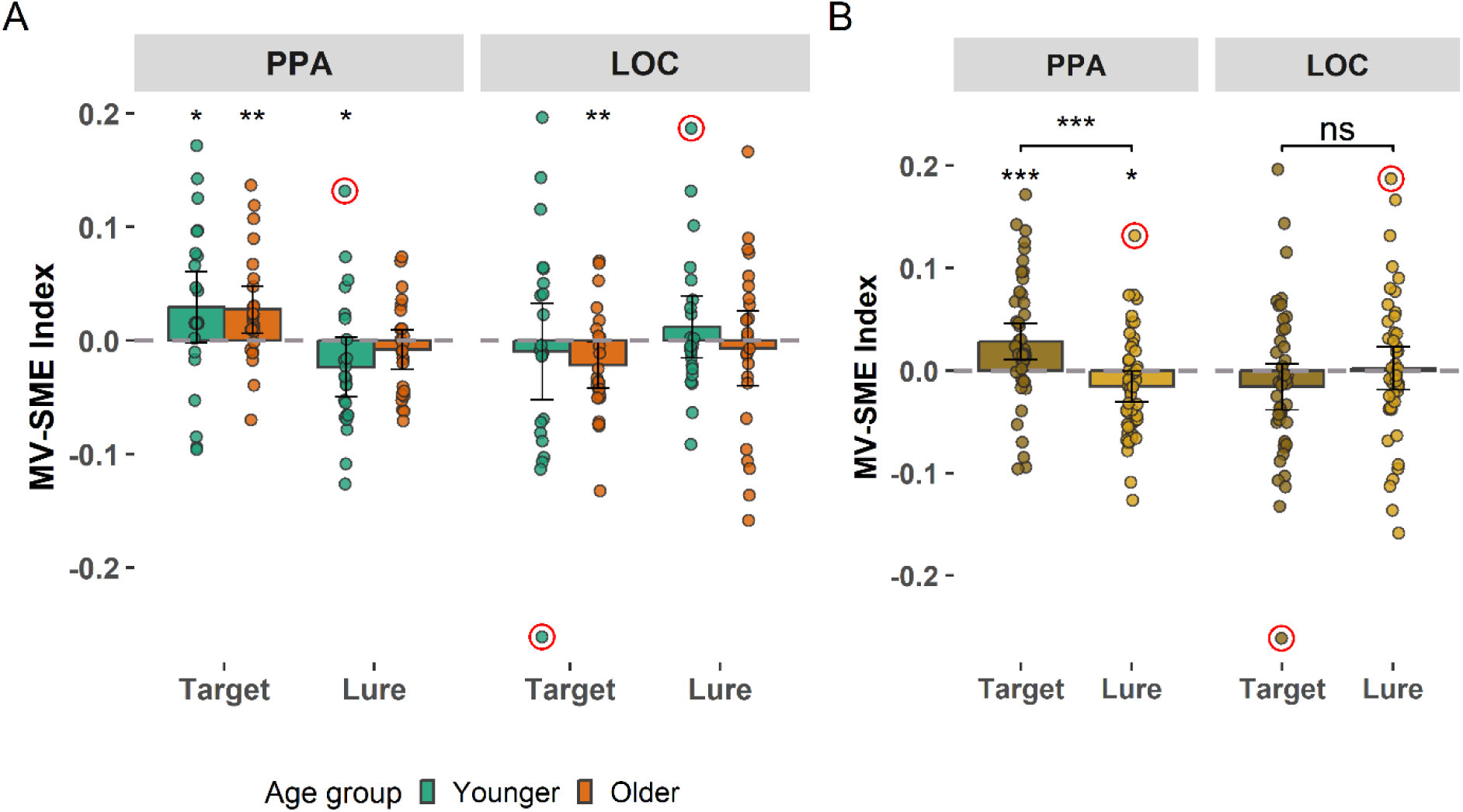
**(A)** Multivoxel SME metrics for younger and older adults, shown separately for scene images in the PPA and object images in the LOC. **(B))** Multivoxel SME metrics collapsed across age group. Error bars represent 95% confidence intervals. Stars above individual bars indicate trial types where one-sample t-tests indicated that the SME index exceeded zero. The brackets indicate the outcomes of two-sample t-tests. (ns p > 0.05, *p < 0.05, **p < 0.01, ***p < 0.001).

##### Outlying data points are circled

To follow up on the ROI × MV-SME metric interaction, separate 2 (target, lure) × 2 (Age Group) mixed-effects ANOVAs were conducted for the PPA and LOC. Consistent with the univariate analysis reported above, age group was retained in the follow-up models despite there being no age-related interaction effects in the omnibus analysis.

In the PPA, results revealed a significant main effect of MV-SME metric, F(1,45) = 32.24, p < .001, partial η² = .417 in the absence of a main effect of age (F(1,45) = .20, p = .655, partial η² < .01), or an MV-SME x age group interaction (p = .254). As is evident from Figure 5, this effect reflected the fact that target MV-SMEs exceeded the MV-SMEs associated with lures, the latter of which demonstrated a negative effect that did not survive multiple comparisons correction. By contrast, no significant main effects or interactions were observed for the object metrics derived from the LOC (min. *p* = .120).

#### 3.2.2. Whole Brain Analysis

As is evident from Figure 6A, the whole brain analysis identified robust SMEs for target scenes in cortical regions that overlapped canonical scene-selective regions (Epstein & Baker, 2019), namely the PPA, Occipital Place Area (OPA), and Medial Place Area (MPA). By contrast, studied scenes that went on to serve as lures and that were correctly identified as such elicited no reliable SMEs. Moreover, for scene stimuli, significant target > lure differences were evident in the same scene-selective regions as those demonstrating target SMEs (see Figure 6B). None of the two-way interaction terms that included age as a factor identified clusters that survived the *a priori* significance threshold.

**Figure 6:**
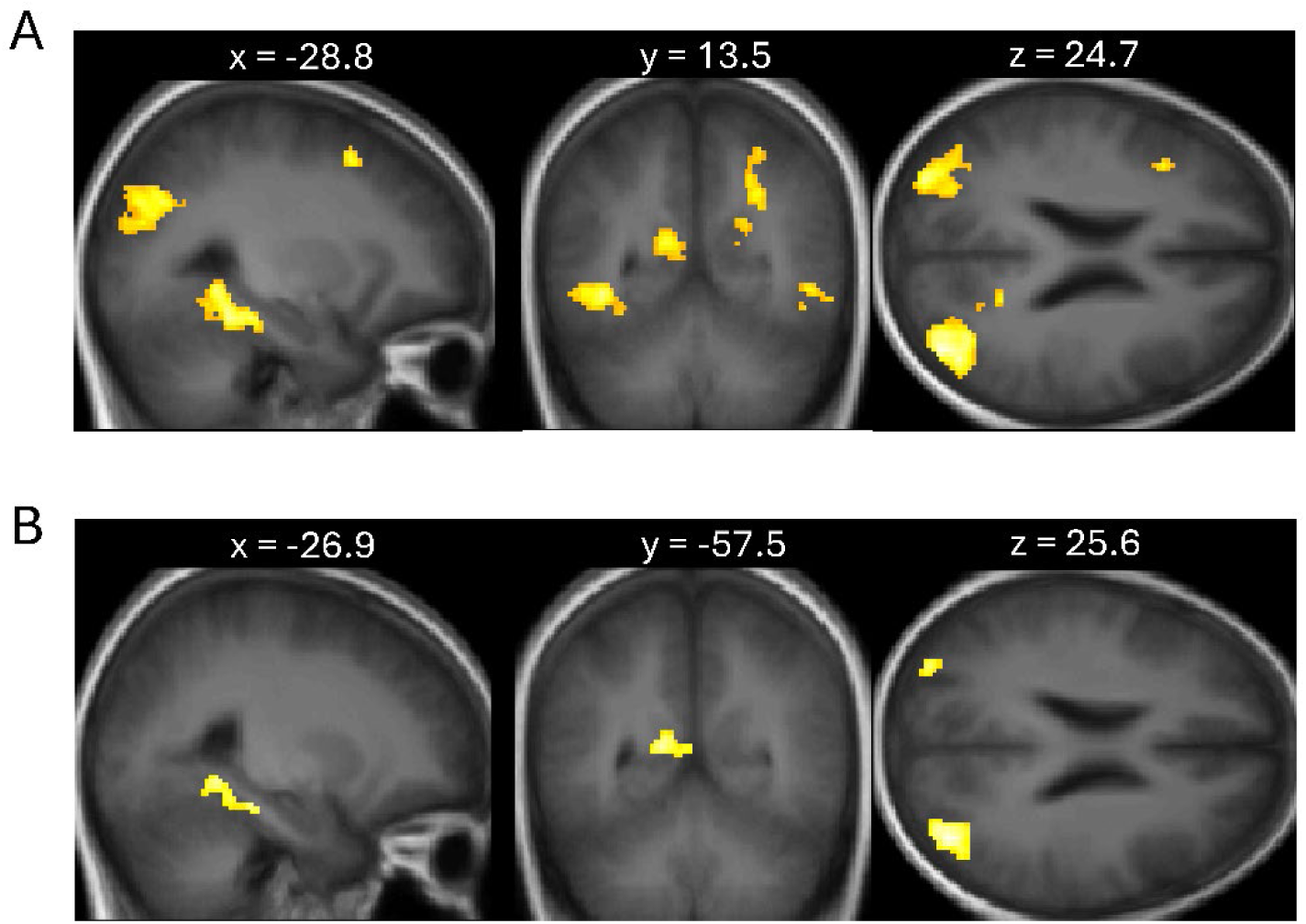
**(A)** Scene Target SME (Scene Target correct > Scene Incorrect) contrast. **(B)** Scene Target-Lure Contrast (Scene Target correct > Scene Lure correct) contrast. The contrasts were plotted on the sample-specific T1 template and were height thresholded at p<.001 with a k = 75 extent threshold.

Thus, consistent with the ROI analyses, the whole-brain analyses identified SMEs for scene targets only, with no evidence of SME × age group interactions. SMEs could not be identified for object images.

### 3.3. Associations between encoding-related neural metrics and mnemonic discrimination performance

As initially reported by Srokova et al. (2024), TLD metrics for scenes and objects were strongly correlated (partial r = 0.670, p < 0.001, controlling for age group), and their separate analysis did not provide additional information. Accordingly, results are reported using the combined TLD metric.

#### Univariate SME metrics and mnemonic discrimination performance

SMEs for targets and lures from each ROI were employed as predictors in four separate models. As described in the Methods, all regression models initially included age group and the interaction between age group and the neural metric. In every model, the interaction terms were non-significant (min. *p* = 0.211) and hence removed from the final model. Following the exclusion of the interaction terms, age group was not a significant predictor in any model (min. *p* = 0.100 for targets; min. *p* = 0.116 for lures). SME magnitude for scene targets was significantly associated with TLD (partial *r*(47) = 0.366, *p* = .012; see Figure 7), whereas scene lure SMEs demonstrated no such association (*p =* .153). Neither object target nor object lure SMEs predicted discrimination performance (min. *p* = .116 for object target SMEs in the LOC).

**Figure 7.**
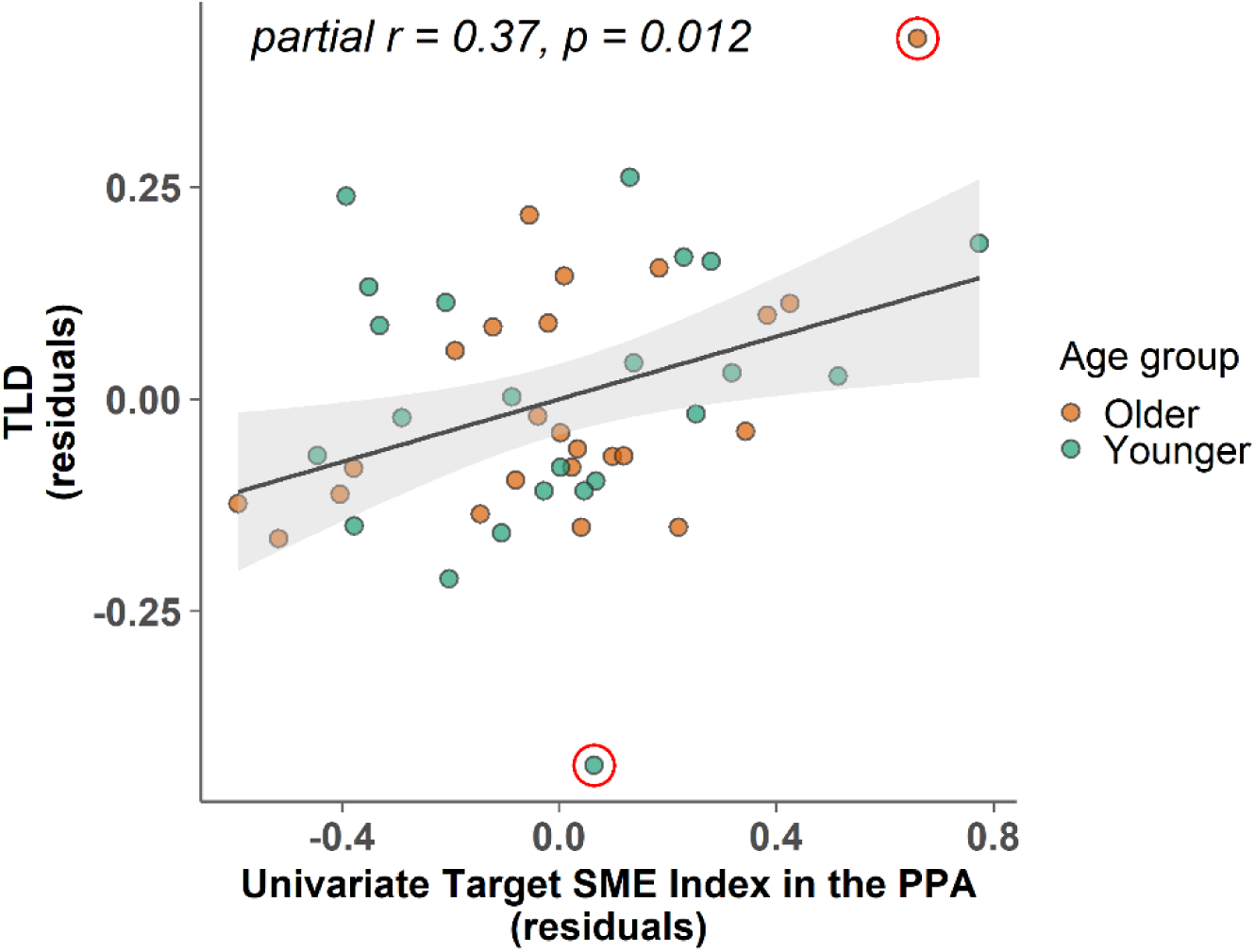
Association between mean TLD and the scene target MV-SME metric in the PPA, controlling for age group. The association remained significant (partial r = 0.32, p = 0.035) after exclusion of the two circled outlying data points.

#### Multivoxel SME metrics and mnemonic discrimination performance

Four separate regression models examined whether the MV-SME index predicted discrimination performance (TLD) within each ROI. Of the four models, including an age group × neural metric interaction term, only the Target LOC MV-SME yielded a significant interaction (*p* = .011).

Follow-up regression analyses conducted separately for the two age groups revealed that Target LOC MV-SME significantly predicted TLD in older adults (*r*(22) = −.51, *p* = .011), but not in younger adults (*p* = .314). However, the association in older adults was driven by a single participant with an outlying score (as defined in section 2.8) on the TLD metric (+2.97 SDs from the group mean). Following the exclusion of the participant, the interaction with the age group and the association for older adults was non-significant (p = 0.538 and p = 0.175, respectively).

For all remaining models, the interaction terms were non-significant (min. *p* = .246) and removed from the final models. Neither PPA (target or lure) nor LOC lure MV-SMEs significantly predicted discrimination performance. The smallest obtained *p*-value was p = .049 (PPA target), which did not survive correction for multiple comparisons. Age was similarly non-significant across all models (min. *p* = .079).

## 4. Discussion

The present study examined whether encoding-related neural activity predicted performance on a Mnemonic Similarity Task (MST) that tested memory for scene and object images. Participants completed a three-choice version of the task, allowing us to separately assess performance and neural activity associated with correct target recognition and correctly identified perceptually similar lures. We employed ROI-based univariate and multivoxel metrics, alongside exploratory whole-brain analyses, to assess whether encoding activity – in the form of SMEs - differentiated correct versus incorrect subsequent target and lure judgments, as well as whether these metrics predicted MST performance.

Two key findings emerged. First, neural activity elicited by scene images, whether indexed by univariate or multivoxel metrics, predicted subsequent memory outcomes for study items that went on to be targets in the subsequent test phase. By contrast, univariate SMEs were evident neither for studied scenes that were replaced at test with a similar lure, nor for any class of object images (for either univariate or multivoxel metrics). Second, univariate scene SMEs demonstrated an age-invariant association with MST performance.

### 4.1. Relevance of findings to models of mnemonic discrimination

A central motivation of the present study was to test predictions derived from the proposal that the ability to correctly identify perceptually similar lure items in an MST depends to a large extent on the recollection of the corresponding studied item (recall-to-reject; e.g., Norman & O’Reilly, 2003; Kim & Yassa, 2013; Yassa & Stark, 2011; Vanderlip et al., 2024; Lee & Stark, 2023). Under this assumption, successful lure identification, like successful target recognition, should benefit from the formation of a high-fidelity memory representation during encoding.

Thus, given that positive subsequent memory effects (SMEs) are typically associated with later recollection-based retrieval (Davachi et al., 2003; Ranganath et al., 2004; Uncapher & Rugg, 2005; 2008), employment of a recall-to-reject strategy should be characterized by qualitatively similar encoding-related activity for study items later associated with correct target recognition and items whose corresponding lures are successfully identified.

The present findings provided little or no support for this prediction. Robust SMEs were observed for studied scenes that were later correctly recognized as targets, but SMEs predictive of the correct identification of scene lures were absent. This dissociation suggests that successful lure judgments did not depend on the encoding or retrieval of the kind of episodic information that supported target recognition. Thus, the current results indicate that successful lure discrimination can depend predominantly on a non-recollective memory signal.

We suggest that these findings are consistent with the proposal that lure identification can be supported exclusively by a familiarity signal. By this view, lures may fail to elicit sufficient familiarity strength to exceed the response criterion for an “old” response but nonetheless elicit a signal sufficiently strong to support a correct ‘similar’ response (cf. Parks et al., 2026). This interpretation contrasts with accounts that emphasize recollective and matching processes as the primary mechanism underlying mnemonic discrimination.

Multivoxel pattern similarity analyses provided converging evidence in support of the foregoing proposals. Consistent with a prior report (e.g., Yu et al. 2025), studied scenes correctly identified as targets exhibited greater pattern similarity than those associated with incorrect judgments, suggesting that the formation of relatively consistent or stereotyped neural representations during encoding supports subsequent retrieval success. In contrast, studied scene images that were replaced with lures did not exhibit this pattern; instead, these images exhibited negative MV-SMEs, albeit only prior to correcting for multiple comparisons.

The present findings arguably fit well with recent theoretical work (Parks et al., 2026) proposing that judgments on similar lures, including those given in three-choice versions of the MST, reflect a mix of three memory signals: recollection rejection (i.e., recall-to-reject), ‘false recollection’, and familiarity. This framework emphasizes that discrimination performance in the MST cannot be attributed to a single mechanism, such as pattern separation, but rather reflects a combination of different mnemonic strategies and processes.

We were unable to identify significant univariate or multi-voxel SMEs for objects. One possible interpretation of these null findings is that object judgments in the present study relied largely on familiarity for both target and similar lure detection (cf. Parks et al., 2026). However, this explanation appears insufficient, at least in respect of objects that went on to become targets, given the numerous prior reports of object-elicited SMEs (e.g., Liu et al, 2021; 2022; Cansino et al., 2015; see Kim, 2011 for review). It is worth noting that a recent study employing PSA to examine the neural correlates of encoding also reported that PSA metrics demonstrated SMEs for scene-like images, but not for images of ‘small’ objects (Yu et al. 2025). Thus, the present null PSA findings appear to have a precedent, arguably making them less anomalous. These observations do not, however, extend to the puzzling null findings for the univariate analyses of object SMEs.

### 4.2. Effects of Age

As discussed in detail by Srokova et al. (2024), age differences in target/lure discrimination were evident only for object images. In parallel with these behavioral findings, there was little evidence for age differences in scene target SMEs (nor indeed, in any other SME metric). These findings are consistent with prior reports that the neural correlates of episodic encoding differ little across much of the adult lifespan (e.g., Brehmer et al., 2016; de Chastelaine et al., 2016; Liu et al., 2020; see Maillet & Rajah, 2014 for review)

### 4.3. Associations between encoding-related neural activity and behavior

Extending prior findings (e.g., de Chastelaine, 2016; Liu et al., 2022), the present results revealed an association between univariate scene SMEs and mnemonic discrimination performance. Specifically, target-related SMEs for scenes predicted TLD scores across participants, suggesting that variability in encoding processes supporting successful target recognition partially explains individual differences in mnemonic discrimination. Importantly, this association was age-invariant. This pattern is reminiscent of prior reports demonstrating age-invariant relationships between scene selectivity in the PPA and memory performance (Koen, 2019; Srokova, 2020; 2024).

### 4.4. Limitations

The present study suffers from several limitations. These include the relatively small sample sizes, which, especially when combined with the low number of trials associated with an incorrect judgment in numerous participants, limited statistical power. Indeed, we cannot rule out the possibility that these limitations contributed to the failure to detect SMEs for the object images. Another limitation is the cross-sectional nature of the study design, which precludes any conclusions about the effects of aging (as opposed to age) on the dependent variables of interest.

## Conclusion

In conclusion, the present study provides evidence that encoding-related activity preceding the administration of an MST can be predictive of subsequent target identification but fail to predict lure identification. Thus, target and lure identification can be supported by distinct (but seemingly age-invariant) encoding operations, suggestive of the engagement of dissociable retrieval processes.

## Acknowledgements

This research was supported by the National Institute of Aging Grants R56AG068149, RF1AG039103, and R01AG082680 as well as BvB Dallas. A.N.Z.A. was supported by the Larry James Family Foundation. S.S. was supported by the Postdoctoral training program of the Arizona Alzheimer’s Consortium, grant T32AG044402. The authors gratefully acknowledge the experimental volunteers for their time and participation, and Joshua Olivier, Nehal Shahanawaz, and Eduardo Hernandez for their assistance with participant recruitment and neuropsychological assessments.

## Author contributions

S.S. and M.D.R. conceptualized the study and developed the methodology. A.N.Z.A. and S.S. conducted the investigation and performed the research. A.N.Z.A. and S.S. carried out the formal analysis. A.N.Z.A. prepared the initial draft of the manuscript. S.S. and M.D.R. contributed to writing, review, and editing. M.D.R. provided supervision and project oversight.

